# Evidences of a component Allee effect for an invasive pathogen: *Hymenoscyphus fraxineus*, the ash dieback agent

**DOI:** 10.1101/2022.10.17.512473

**Authors:** Simon Laubray, Marc Buée, Benoît Marçais

## Abstract

Invasive pathogens are a major threat to forest health especially in managed forest with low diversity. The die-back of European *Fraxinus spp*. caused by the fungus *Hymenoscyphus fraxineus* is the latest example of pathogen invasion causing widespread damage. Host resistance and environment, in particular stand factors were shown to strongly impact disease severity on European ash. The fact that *H. fraxineus* reproduce mostly through heterothallic sexual reproduction suggest that an Allee effect could limit the mating success at low host densities, thus limiting inoculum production and disease development. Populations of *H. fraxineus* were monitored during the fruiting period in a network of stands across a host density gradient in forest and non-forest environment. Ash dieback, basal area of ash, density of infected ash leaf debris (rachis) and apothecia in the litter and ascospores load in the air were determined in the different environments during two years. We showed significant differences between forest and non-forest environment with ash dieback, infection rate and inoculum production higher in forest settings. Host density significantly affected disease development, with crown dieback, density of infected rachis in the litter and inoculum production increasing with host density. We also demonstrated that fruiting rate, i.e. the number of apothecia per infected rachis dry weight, is strongly dependent on infected rachis density. Inoculum production is therefore limited at low host densities. Such a component Allee effect could be important in *H. fraxineus* epidemiology and invasion dynamic.

## Introduction

Emerging infectious diseases are major threat to forest ecosystems. They are frequently induced by invasive fungi and can have very important consequences on the biodiversity (Desprez-Loustau et al. 2007, 2009). The most frequently cited examples of tree species that were drastically reduced by an invasive pathogen include chestnut in USA with *Cryphonectria parasitica* or elm in Europe and the USA with *O. novo-ulmi*. The rate of invasive forest pathogens arrival has been increasing from 1980 to 2008 in Europe, with a particular importance of ascomycetes (Santini et al. 2013). Invasive pathogens currently represent a proportion of about 50% of the disease cases reported by the forest health survey system in France, with a high share of currently severe epidemics such as ash dieback caused by *Hymenoscyphus fraxineus* (Desprez-Loustau et al. 2016).

As any alien species, invasion success of exotic fungal pathogens depends on overcoming different barriers (Blackburn et al. 2011) to thrive in their new environment. The pathogen needs to be transported to the new environment, then it must adapt to the environment, which implies finding available hosts (Engering et al. 2013). During this establishment step, the invasive pathogen needs to reproduce and to survive efficiently off season to produce a stable population. The next step is the spread throughout the new territory that depends on the dispersal abilities and on the availability of favourable habitats. The establishment is a critical phase; it depends strongly on the habitat suitability, but also on the pathogen population dynamic (Taylor and Hastings 2005) and on the host density present in the landscape (Park et al. 2001; Condeso and Meentemeyer 2007). Founding populations of invasive species are often small and could be subjected to an Allee effect. Allee effect is a mechanism that may reduce the population growth at a low density and thus reduce the establishment likelihood. Stephens et al. (1999) defined the Allee effect as “a positive relationship between any component of individual fitness and either numbers or density of conspecifics”. He further distinguishes the component Allee effect, which affects an individual fitness component and demographic Allee effect which affects the total species fitness. A component Allee effect would be a positive correlation between a fitness parameter such as the mating success with the population density whereas the demographic Allee effect implies a relationship between the per capita growth rate and the population density and is more difficult to demonstrate. An Allee effect may limit the population growth rate at low density with different strength. Strong Allee effect result in negative growth rate at very low population density, leading to extinction while weak Allee effect will only reduce the population growth rate through density dependence. In invasive species, the Allee effect may result in a latency phase at the colonisation front in which population growth is highly dependent on population density followed by an important growth increase when the population reach a so-called Allee threshold (Hastings 1996; Veit and Lewis, 1996). In populations that are denser than the Allee threshold population density is no longer the limiting factor for growth. In addition, another factor limiting invasion is the strong dependence of the invasive pathogen population on the population density of its host. A high host abundance in the landscape allows the pathogen to colonize the area more easily (Jules et al. 2002). Moreover, as other parasites, plant pathogens are known to be a driver of host population size (Cobb et al. 2012). Pathogens appear to strongly structure host populations limiting growth and regeneration or induce mortality. On the contrary, low host density could limit the development of the pathogen population especially if its dynamic is subject to Allee effect.

The invasive pathogen *H. fraxineus* was described as the responsible of ash dieback (Kowalski et al 2009; Gross et al. 2014a; Baral et al 2014). First observations of European ash dieback were noted in Poland in the 90’s (Przybył 2002) and was attributed *a posteriori* to the introduction of the ascomycete *H. fraxineus* (Gross et al. 2014a). This pathogenic fungus of common ash (*Fraxinus excelsior*) and narrow-leaved ash (*Fraxinus angustifolia*) is native to East Asia (Zhao et al. 2013; Gross et al. 2014b; McMullan et al. 2018) and spread through Europe to reach Ireland in 2012 (Short et al. 2019), Montenegro in 2016 (Milenković et al. 2017), north of Spain in 2021 (Stroheker et al. 2021). This pathogen caused serious damage in the ash stand, threatening associated species and biodiversity (Pautasso et al. 2013). *H. fraxineus* biological traits and the constant spread rate observed over time in France led Hamelin et al (2016) to suggest the existence of a component Allee effect caused by the mating success.

This hypothesis is based on three facts. First, *H. fraxineus* is a heterothallic fungus, *i*.*e*. successful sexual reproduction needs presence of two mating types (Gross et al. 2012). The reproduction of this foliar pathogen occurs early in summer on residual leaf debris present in the litter (rachis= petiole + central vein of the compound leave) and result in formation of apothecia. Despite rare observations of apothecia production on other infected tissues by *H. fraxineus* (Kirisits and Freinschlag, 2012; Kowalski and Holdenrieder, 2009; Wylder et al. 2018), the infected rachis density in the litter could be considered as main sexual reproductive population. Moreover, the ascospores released by apothecia are the main inoculum vector. Indeed, conidia produced on the rachis are believed to act only as spermatia (Gross et al. 2012), and their spread by splashing is limited to few centimetres or few metres. As a consequence, fecundation might be limited at low density of infected rachis in the litter.

Second, the spread of the pathogen is strongly linked to the host density. *Fraxinus* is widely present in the landscape and its distribution area extends from South of Scandinavia from North of Turkey and North of Spain (*F. excelsior*) and South of Europe and Turkey (*F. angustifolia*) (EUFORGEN) making it easier for the pathogen to spread. A study showed the importance of ash density and tree cover fragmentation for establishment and disease development at the landscape scale (Grosdidier et al. 2020). Damage on the crown and collar was severe in dense ash stands in forest while isolated ashes out the forest were less affected. Moreover, the severity of ash dieback is correlated with the load of inoculum produced in the litter (Marçais et al. 2016). A component Allee effect on the reproduction success could further reduce the inoculum production at low ash density.

Finally, the environmental conditions strongly constraint the severity of ash dieback. The inoculum production is particularly affected by the level of ambient humidity. Abundant apothecia formation needs high humidity (Hietala et al. 2013; Dvorak et al. 2016; Grosdidier et al. 2020) and thus, factors influencing ambient humidity such as tree cover, vicinity to river, topography and precipitation will impact inoculum production and damages to the ashes (Havrdová et al. 2017.; Enderle et al. 2019; Skovsgaard et al. 2017; Grosdidier et al. 2020). Further, temperatures higher than 35°C are lethal for *H. fraxineus* which explains limited spread in the Mediterranean region with hot summers (Grosdidier et al. 2018; Hauptman et al. 2013).

The objective of this study was to unravel the relationship between host density and their leaf litter production (rachis density), their crown health status, infected rachis density in the litter as reproductive pathogen population size and inoculum production by apothecia density in ash dieback. Assuming that the population size of *H. fraxineus* can be measured by infected rachis density in the litter and depends on ash density and level of infection, we hypothesised that a component Allee effect on the success mating limits inoculum production and the ash dieback at low population density (host and pathogen). We monitored the population dynamics of *H. fraxineus* during the period of inoculum production in an ash density gradient in forest and open landscape.

## Materials and methods

### Stand Characterisation

A network consisting in 20 plots in forest stands and 10 plots in hedges and small woods was installed on the village of Champenoux in North East France (WGS84 48.7521N 6.3409 E, Fig. 1). The network was established in order to obtain a gradient of host density. Host density was measured by the basal area of ash (*Fraxinus excelsior*). At each studied location, three concentric circular plots were established. All tree stems with a diameter at breast height (DBH) over 7.5cm were measured within a radius of 7 m (154 m) while only tree stems with a DBH over 22.5 cm were measured in a radius of 7-16 m (805 m^2^) and with a DBH over 47.5 cm in a radius of 16-21 m (1386 m^2^). The basal area of ash in m^2^. ha^−1^ was computed by summing individual stem area at breast height weighted by the sampling surface. We also measured the basal area of other trees species present in the stand and total basal area was used as a proxy of canopy closure. Ash basal area was used to explain local density of rachis in the litter. However, in order to evaluate the impact of ash density on ash health, we weighted basal area by the rate of tree cover within the 100-m radius around the points; this enabled to account for the very patchy presence of trees outside forest stands. The tree cover rate was computed with QGIS software using an IGN shape file corrected with aerial photograph (BD Ortho^®^ edition 2018 and BD FORET^®^ version 2.0 available in Web Map Service flow on https://geoservices.ign.fr). The size and health status of ash trees included in the basal area assessment was recorded with their DBH measure and the following rating of crown mortality: 0-10% (healthy), 10-50% (symptomatic), 50% -75% (declining) and >75%. The health status of an ash stand was computed as the mean of the tree ratings (using the median of their health class) and the ash size was estimated with the mean of DBH. Four plots were moved by a short distance in 2021 because the 2020 location was compromised by logging. The meteorological conditions covering the sampling periods (from 1 June to 31 July 2020 and 2021) were collected at the Champenoux weather station (Figure 1). The heat level was expressed by the mean of daily maximal temperature and mean of daily mean temperature, the humidity level was expressed by mean of daily humidity and sum of precipitation.

**Fig. 1.**
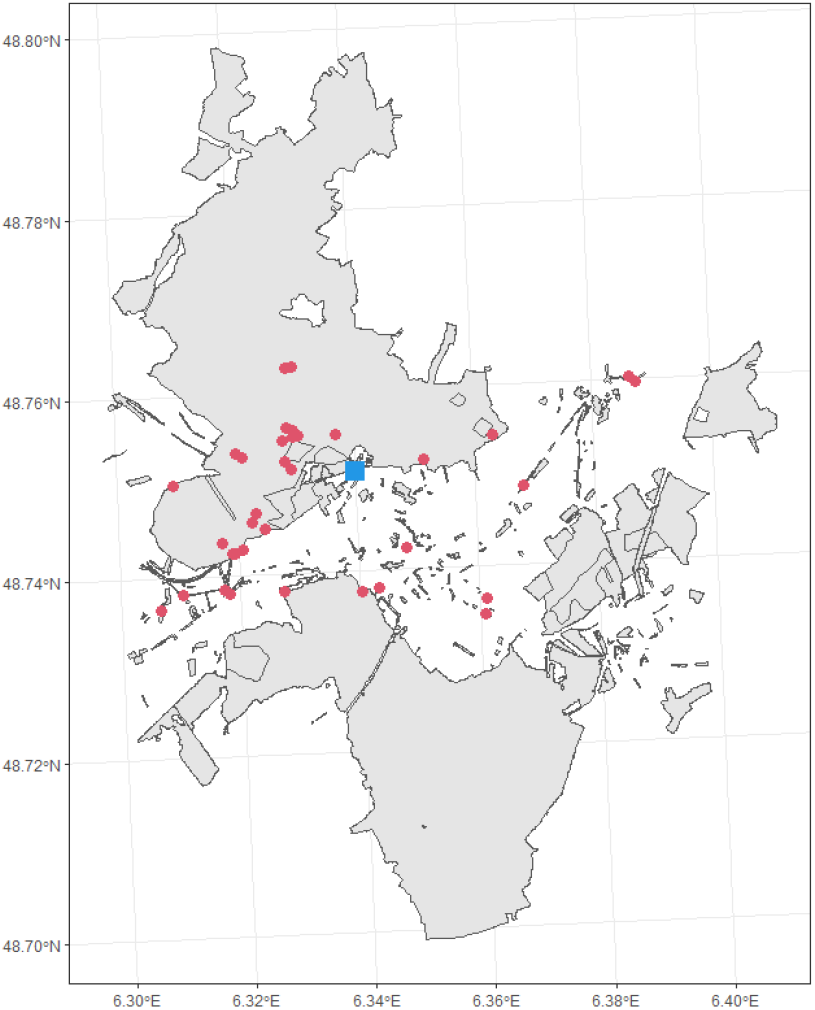
Plot distribution around Champenoux village of ash stand sampled for ash density, health status, rachises densities (total and infected), apothecia density and ascospores trapping (red point). Meteorological station (blue square). Wooded area appears in grey on the map.

### *Hymenoscyphus fraxineus* population size and inoculum production

In each stand, the density of ash rachises at the soil surface and their frequency of colonisation by *H. fraxineus* was determined in June 2020, June 2021 and July 2021. For that, all ash rachises present along ten 0.1-m^2^ areas located along two 10-m perpendicular transects were collected (10-cm wide area along the transect on each other meter). The rachises were sorted in the laboratory according to their colonisation status by *H. fraxineus*. Rachis with the presence of a distinct black pseudosclerotial plate is characteristic of *H. fraxineus* infection while rachis un-colonized by the pathogen remain light brown to grey with absence of a pseudosclerotial plate and are considered as healthy. To confirm the assessment, a part of rachis identified as infected or healthy were placed in moist chamber for 8 weeks to monitor appearance of apothecia. The proportion of rachises from both categories that produced *H. fraxineus* apothecia was then computed.

Infected and healthy rachises collected on plots and sorted in lab were dried for 48h at 50°C and then weighted. The number of apothecia present on these rachises was counted during the sample collection of rachises. The mean dry weight of rachis (infected and total) and the mean apothecia frequency per plot were then computed in g.m^−2^ and No. of units.m^−2^, respectively. The fructification rate was computed as the number of apothecia per infected rachis dry weight (N.g^−1^). Data on number of apothecia and infected rachises density per m^2^ from previous work were used in the analysis of fructification rate (15 plots sampled in 2012 from Grosdidier et al. 2018, 23 plots sampled in 2016 and 31 plots sampled in 2017 from Grosdidier et al. 2020). The sampling method is similar to our method except that the infected rachises density is measured in length of rachis per unit surface (cm.m^−2^)., the rachis density was converted in weight per m^2^ (g.m^−2^) according to the relationship (L = 163.73 * DW, where L is the length in cm and DW, the dry weights of rachises, Grosdidier et al. 2020).

The spore trapping method developed by (Grosdidier et al. 2017) was used to determine the air load of *H. fraxineus* ascospores on the studied plots. Shortly, the spore traps are passive traps composed of cellulose filter (Whatman™ 150 mm diameter Cat No 1001-150) imbibed of 5 ml of glycerine and place on a styrifoam block at 1 meter above the ground. Three spore traps were set up per plot and left exposed for two consecutive periods of 15 days (22 June to 8 July and from 9 to 22 July). After exposure on the plots, the filters were recovered and put individually in plastic bags. In the laboratory, 30 ml of 4x TE buffer (40 mM Tris-HCl, 4 mM EDTA, pH 8.0) heated at 60°C was added into each plastic bag with filter. The filters were gently hand rubbed through the plastic to separate the captured particles from the filter. The TE buffer was then transferred in 50 ml vials and centrifuged 15 min at 2700 g. The supernatant was removed to keep approximately the bottom 3 ml of suspension containing most of the particles. This 3 ml of suspension was transferred in two 2-ml microtubes, centrifugated 5 min at 18 620 g and, 750 μl of supernatant was removed from each tube. The remaining 750 μl were vortexed, pooled in one tube and centrifugated once again 5 min at 18 620 g. The 200 μl bottom of the concentrated particles solution was kept at -20°C until DNA extraction.

DNA was extracted from the 200 μl concentrated particles solutions using the DNeasy plant mini kit (Qiagen). Two 3-mm and twenty 2-mm glass beads were added to the particles solutions together with 400 μL of lysis buffer and 4 μl RNase. The samples were grounded twice 50 s at 6 m.s^−1^ with FastPrep-24 MP BIO and incubated 30 min at 65°C to lyse the cell. The following steps were done as described by the manufacturer. Total DNA was then eluted in 200 μl AE buffer.

The number of *H. fraxineus* ascospores from spore traps was quantified by qPCR using the method developed by Ioos et al. (2009). The 15 μl of reactional mix were composed of 1x Brillant II qPCR master mix (Agilent Technologies), 0.03 μM ref dye provided with the master mix, 0.01 U.μl^−1^ UDG (New England BioLabs), 0.3 μM each Cfrax primers (Cfrax-F, 5′-ATTATATTGTTGCTTTAGCAGGTC-3’ and Cfrax-R, 5′-TCCTCTAGCAGGCACAGTC-3′), 0.1 μM Cfrax-probes (Cfrax-P, 5′-FAM-CTCTGGGCGTCGGCCTCG-BHQ1-3′) and 2 μl template DNA. The real time reaction was performed in a Quantstudio 6 thermocycler (Applied Biosystem). The qPCR reaction was initiated by first pre-cycling step at 37°C for 10 min for UDG activation and the initial denaturation step at 95°C for 15 min followed by 50 cycles of denaturation at 95°C for 15 sec and hybridization /elongation at 65°C for 55 sec. The ascospore quantification was performed using ascospore solutions obtained by tenfold cascade dilution with from 50 000 to 5 ascospores per μl.

### Statistical analyses

Crown decline, rachis densities in the litter (total and infected by *H. fraxineus*) and the infection rate of rachis in the litter were analysed with generalised linear mixed models (glmm) using the R library glmmTMB. In both cases, the plot was declared as random effect.

The crown decline rate and the rachis infection rate were modelled using a Beta-distribution that is well adapted to variables lying between 0 and 1 (Figueroa-Zúñiga, et al 2013); we used the logit link function. The explicative variables were the host density (ash basal area) and environmental variables, with the measured year added for the rachis infection rate as fixed factor. The densities of ash rachis in the litter which is a positive continuous variable was modelled with the Gamma distribution with a log link function. Ash density, crown decline and mean ash diameter were included as explicative variables. The apothecia density, the amount of ascospore detected in the spore traps and the infected rachis density were modelled with the negative binomial using an identity link function. The sampled plots were declared as random factor.

These GLMs relationships were used to build a Structural Equation Modelling (SEM) using the R-package “piece-wise” (Fig. 3). SEM highlighted the correlation between host variables (ash basal area, average size and health status, total rachis density), *H. fraxineus* population variable (infected rachis density) and inoculum production (apothecia density and amount of ascospores). The year of sampling was added to evaluated the summer variability and the environment to compare the forest to hedge and small wood. The coefficient estimates were standardised in order to compare the effect of the different parameters.

To assess whether a component Allee effect was present for mating success, the relation between the apothecia production rate and *H. fraxineus* population size (infected rachis density in the litter) was studied. The fruiting rate (***τ***) was define as the number of apothecia produced per unit of dry weight of infected rachis present in the soil litter:

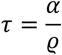

Where ***α***, is the apothecia number per m^2^ and ***ϱ***the infected rachis density (g.m^−2^). Without Allee effect, ***τ*** is constant, while in presence of a component Allee effect, it should be positively correlated with the infected rachis density. The Allee effect should be most effective at low rachis density, with no density dependence above a threshold value (Allee threshold). Therefore, the relationship between ***τ*** and the infected rachis density can be modelled with a Gompertz function (equation 1) with the hypothesis that the fruiting rate may be zero at very low density, positively dependent in low density and reach an optimum at high density superior of Allee threshold (Fig.2).

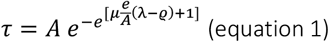

**Fig. 2.**
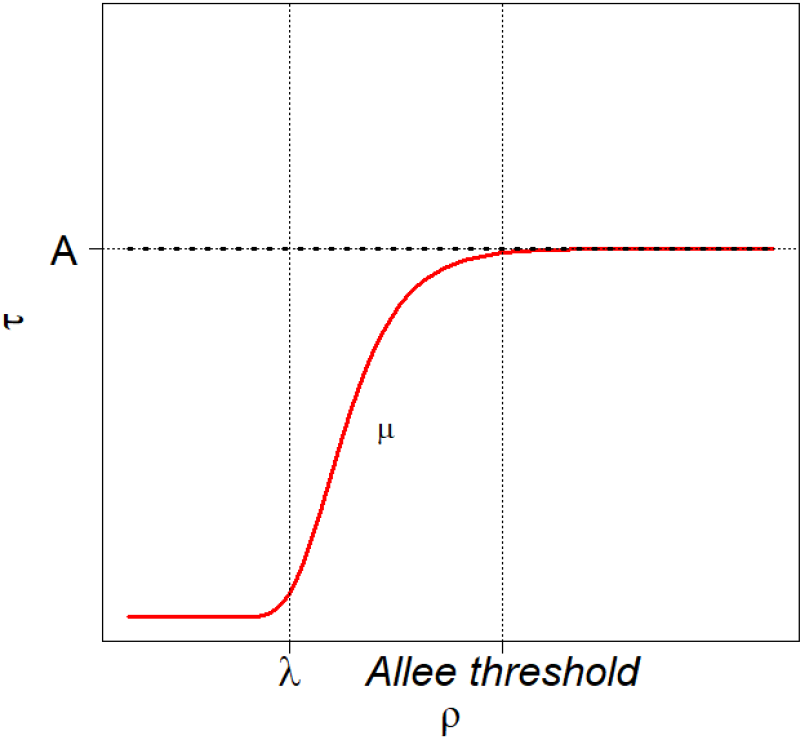
Theoric curve of (**ϱ**) with in bold dashed black line no Allee effect and in red line presence of Allee effect according to Gompertz equation. With the parameter of Gompertz equation: **A**. as the average **τ**. reached for density **ϱ**. > Allee threshold, **μ**. as the maximal slope and **λ**. the minimum value of density **ϱ**. where **τ**. > **0**

With ***A***, the maximum fruiting rate (optimal mate encounter), ***μ***, the maximum slope and ***λ***, the strength of Allee effect, i.e the value of infected rachis density under which mate encounter does not occur and no apothecia are produced. The ***Allee threshold*** is defined as the value of infected rachis density ***ϱ*** above which the fruiting rate ***τ*** is ***A*** on average and does not depend on ***ϱ*** anymore (Fig. 2).

The Gompertz function was fitted in a Bayesian framework using the R2jags package. The fructification rate τ was assumed to follow a Gamma distribution with the mean following equation 1. Parameters ***A*** and ***μ*** were assumed to depend on environment (forest or hedge/small wood) and on a year random effect. Flat priors were assumed for the parameters: normal distribution N (0,0.0001) for parameters for ***A***, uniform distribution U (20,100) for parameters for ***μ***, uniform distribution U (0,5) for ***λ*** and uniform distribution U (0,100) for variance of the random factors and of the Gamma distribution. We run 3 MCMC chains for 100000 iteration with a burn-in of 75000 and a thin of 10. The convergence was assessed by Gelman-Rubin tests. The fit of the model was checked by comparing the mean and dispersion of observed data and of data simulated according to the model. We tested whether a significant Allee effect was present by computing the value of the **Allee threshold**, defined as the value of infected rachis density ***ϱ*** for which average fructification rate ***τ*** reached 0.99 **\* *A***, and assessing whether this threshold was significantly different from 0.

The total density of rachises in the litter needed to reach the Allee threshold depends on the infection rate (equation 2). On the other hand, the log of total density of rachises in the litter is also proportional to ash basal area (equation 3).

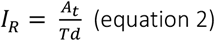

and

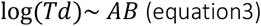

Where ***I***_***r***_ the litter rachises infection rate minimum to reach Allee threshold is determined by ***A***_***t***_ the infected rachis density of the Allee threshold divided by ***T***_***d***_ the total rachis density provided by ***AB***, the ash basal area

We estimated the litter rachises infection rate needed to reach the Allee threshold depending on the ash basal area AB. For that, a bootstrap procedure was used. Values for the Allee threshold were derived from the Bayesian analysis of *H. fraxineus* fruiting rate. The total ash rachis density (infected and healthy by *H. fraxineus*) was estimated depending on the ash basal area according to the Gamma fitted regression. First, 7500 set of simulated model parameters were generated assuming a multinormal distribution of the parameters using the function mvr-norm of the MASS R package. Using this set of simulated parameters and average values of ash dieback rating and DBH, the total rachis density was computed for plots with an ash basal area AB from 1 to 40, and ash dieback and DBH values equal to the mean values of the studied plots. The litter rachises infection rate needed to reach the Allee threshold was then computed as the ratio between the Allee threshold and the total rachis density. We computed its mean and its 2.5% and 97.5 quantiles.

## Results

The ash basal area gradient extends in forest from 2.2 m^2^.ha^−1^ to 18.3 m^2^.ha^−1^ with 50% of stands below 4.6 m^2^.ha^−1^. In hedge and small wood, this basal area is between 2.5 m^2^.ha^−1^ and 37.5 m^2^.ha-1 with a median of 10.7 m^2^.ha-1. Pure ash stands were no longer present in forests around Champenoux. The highest ash basal areas were observed in small wood, where ashes were concentrated in isolated small areas. The ash basal area obtained after weighting by tree cover in a 100-m radius in hedge and small wood range between 0.12 m^2^.ha^−1^ and 13.5 m^2^.ha^−1^ with a median at 1.8 m^2^.ha-1. The weighted range of total basal area was between 0.12 m^2^.ha^−1^ and 35.8 m^2^.ha^−1^ with a median of 19.7 m^2^.ha^−1^.

Weather conditions of the summer 2020 and 2021, during the apothecia production and ascospores release period (June and July), were relatively different. While average daily maximal temperature was similar (p>0.05), air humidity and precipitation were very different, with summer 2020 being drier than summer 2021 (p<0.001) (Table 1).

**Table 1.** Weather conditions during the apothecia production (June and July). Max.Temp : average daily maximal temperature, Moy.temp: average daily temperature, Moy.Hum: average air humidity, Sum.Prec: precipitations sum.

| Year | Max.Temp | Moy.Temp | Moy.Hum | Sum.Prec |
| --- | --- | --- | --- | --- |
| 2020 | 24.6±1.1°C | 18.4±0.7°C | 65.9±3.3%*** | 76.0mm |
| 2021 | 24.3±0.9°C | 18.6±0.6°C | 79.7±2.8%*** | 221.5mm |

The SEM analysis showed strong positive relationships between the population dynamic of *H. fraxineus* and the density of its host (Fig. 3). The amount of ascospores increased with the apothecia density (0.001, p<0.05), which in turn increased with the infected rachis density present in the stand litter (0.017, p<0.001). The infected rachis density depended on the total rachis density (0.26, p<0.001) which itself depended on ash basal area (0.07, p<0.01) and ash diameter (0.03, p<0.01). Increasing crown dieback had a negative effect on total rachis density (−0.05, p<0.01). Crown dieback was more severe when ash density, i.e. ash basal area, was high (0.88, p<0.05). Thus, in dense ash stands with severe dieback, the total rachis density was reduced which could decrease the density of infected rachis. The apothecia density and the amount of ascospores captured were affected by the environment with lower values in hedges and small woods (respectively 0.01, p<0.001 and 0.1, p<0.001). The year affected the apothecia density and even more the infected rachis density (0.57, p<0.05), with lower value in 2020.

**Fig. 3.**
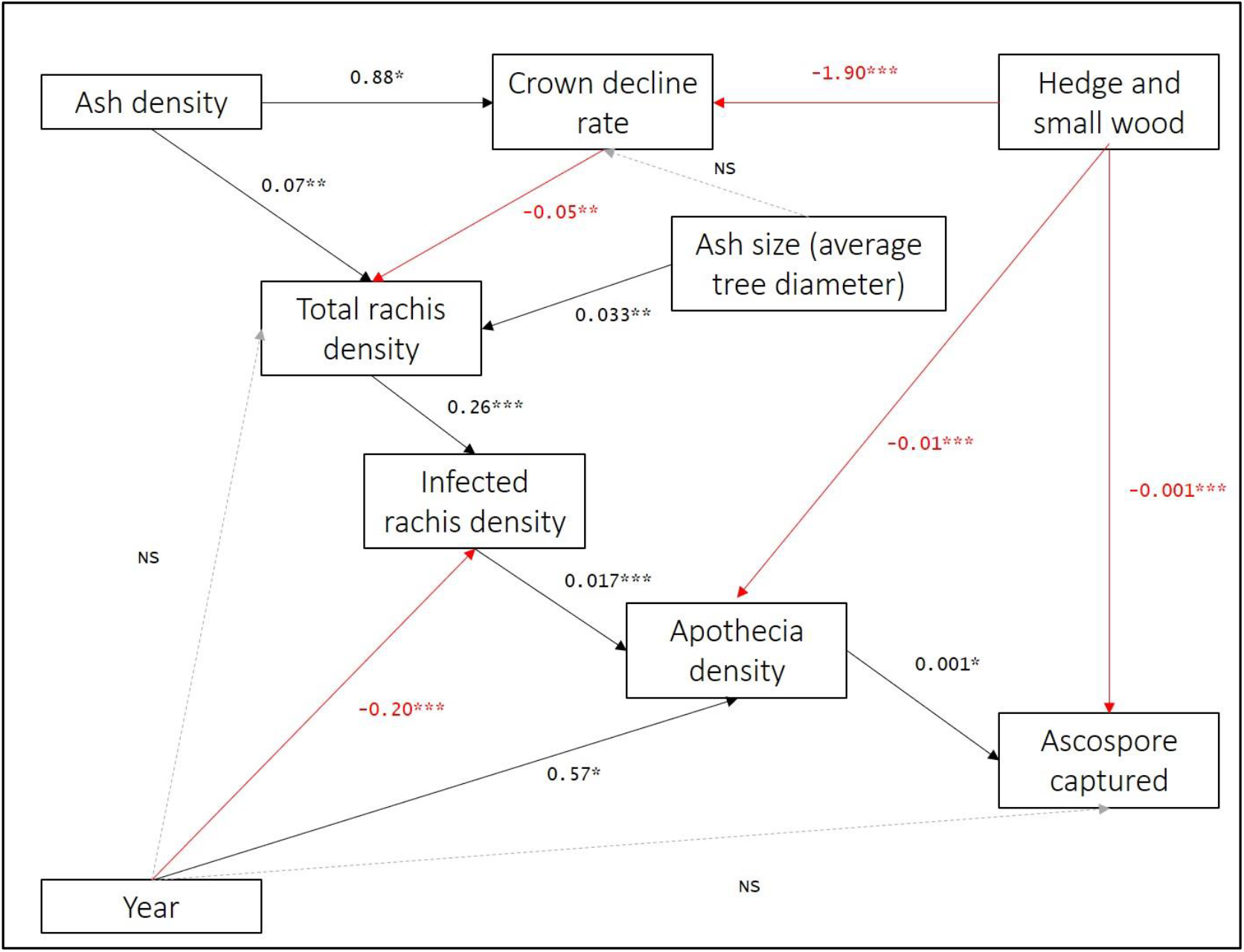
Structural equation modeling (SEM) to highlight the relation between hosts parameters (ash density and total rachis density), tree health status (crown decline rate), pathogen population (rachis density infected H. fraxineus) and fungal inoculum production (apothecia density and captured ascospores) according to year and environment. Black line positive correlations, red line negative correlations and dotted line no correlations, the coefficients of correlation estimated were scaled with * p <0.05, ** p <0.01 and *** p <0.001.

### Crown decline and ashes density

A total of 302 ashes was assessed for crown dieback, with 30% of trees located in forest plots and 70% in hedge and small wood plots. The ash trees were overall healthy with 128 trees that rated as 0.05, 120 trees as 0.3 and the most severe dieback classes unfrequently observed (36 trees rated as 0.625 and 18 trees as 0.825, Fig. 4a). Health status, i.e. mean plot crown decline, significantly deteriorates with increasing host density (0.14 ± 0.01, p<0.001) (Fig. 4b). Healthy ashes were predominant at ash density less than 2 and their number decreased as ash density increased. By contrast, the number of symptomatic and declining trees increased with ash density. Additionally, the mean crown decline was greater in forest than hedge and small wood with a mean crown decline of 0.41±0.05 (IC95%) in forest and 0.19±0.03 (IC95%) in hedge and small wood (Fig. 4b p<0.001). Ash trunk diameter was not related to dieback severity (Fig. 4c, p>0.05) and did not differ between forest and hedge/small wood.

**Fig. 4.**
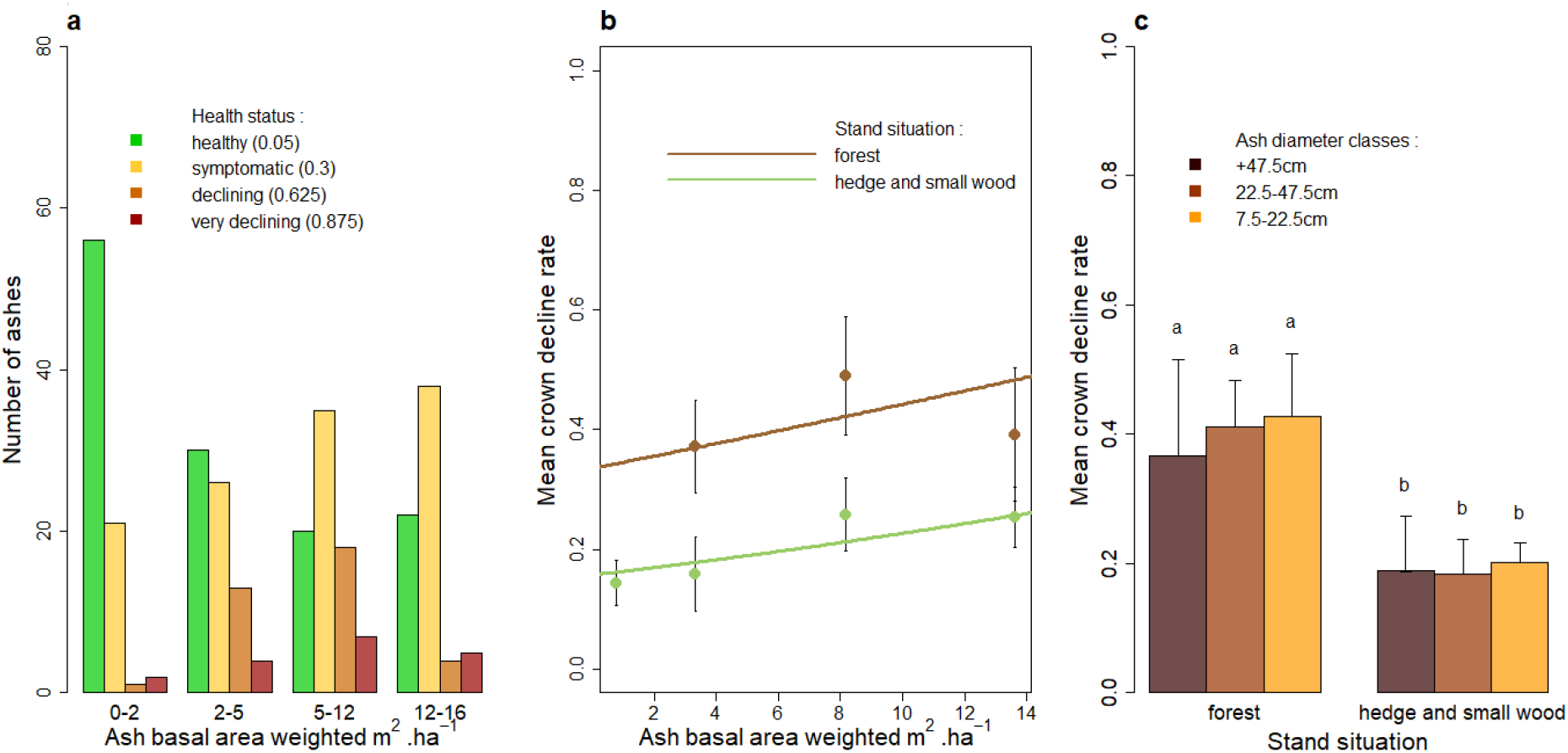
**a** Ash distribution according to ash density and health status class (with median crown decline rate for each class). **b** Mean crown decline rate of the ash stand according to ash density and the environment. **c** Mean crown decline rate in different ash diameter classes in forest and hedge and small wood.

### *Hymenoscyphus fraxineus* population size and inoculum production

The fruiting test confirmed that the classification of rachises as infected / healthy was adequate. A total of 3.6 ± 1.1 % of rachises classified as healthy produced *H. fraxineus* apothecia, while 96.0 ± 1.2 % of rachises classified as infected were producers.

Total rachis density present in the litter significantly depended on ash density, mean ash diameter and health status of ash crowns (Fig. 5). The total rachis dry weight per square meter significantly increased with the ash basal area and mean ash diameter (respective coefficients 0.07 ± 0.01, p<0.001 and 0.02 ± 0.01, p<0.01). The deterioration of ashes crowns reduced the density of rachis present in the litter (−1.25 ± 0.55, p<0.05) (Fig. 5a). The density of rachis per unit ash basal area was decreased with increasing mean crown dieback rate (−1.64 ± 0.60, p<0.01) (Fig. 5b). In hedge and small wood, the rachis production was slightly lower than in forest situation (−0.54 ± 0.24, p< 0.05) (Fig. 5b). The rachis infection rate showed no relationship with the ash density (p>0.05). The mean rate of infected rachis was higher in 2020 (0.55 ± 0.07) than in 2021 (0.26 ± 0.05) (p<0.001 Fig. 5c). However, the infection rate was similar for plots in forest or in hedge / small wood (p>0.05, Fig 5c).

**Fig. 5.**
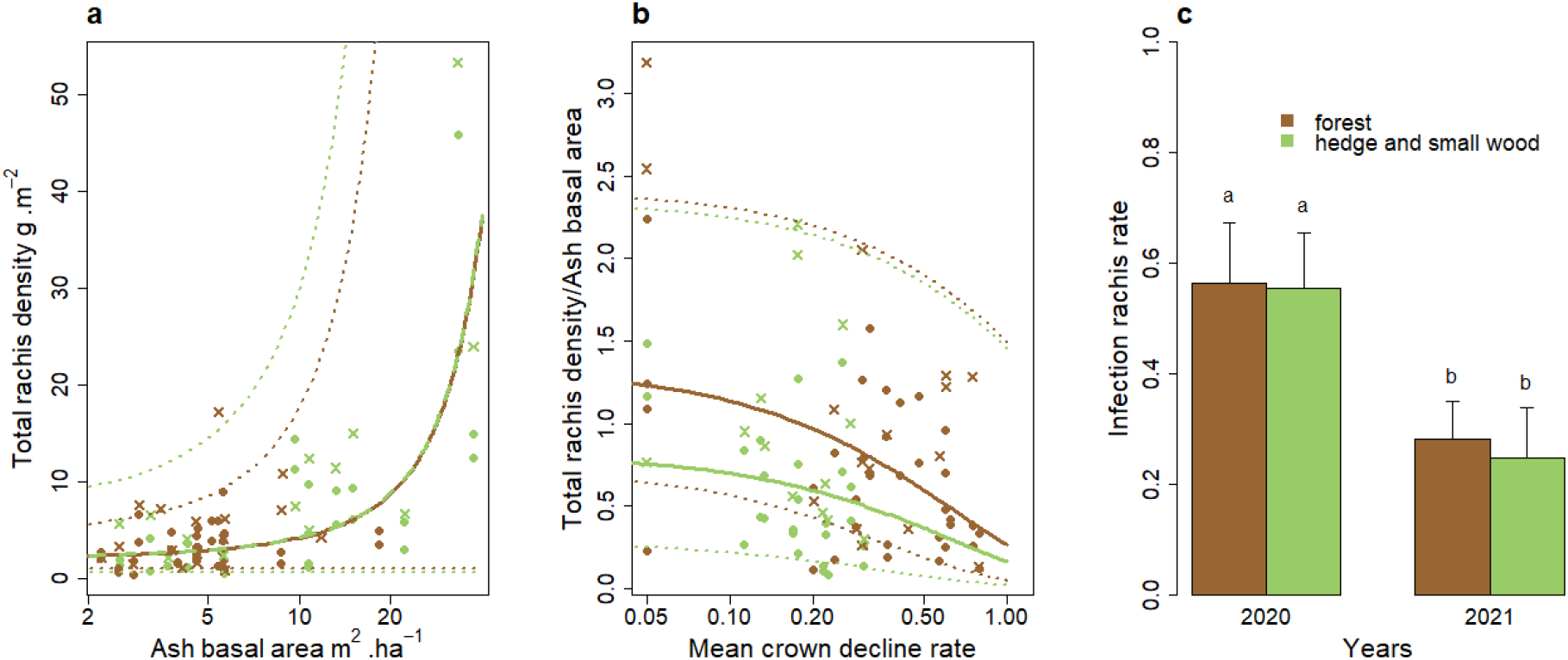
**a** Total ash rachis density collected in the litter in forest (brown points) and in hedge and small wood (green points) according to ash density and their associated total rachis density, confidence intervals at 97.5% (dotted lines). **b** Rachis production by ashes according to mean crown decline rate for forest situation (brown), hedge and small wood (green) and the fitted regression model(line) and confidence intervals at 97.5% (dotted lines). **c** Infection rate according to ash density for forest situation (brown), and hedge and small wood (green), for year 2020.

In 2020, the amounts of apothecia produced were very weak. No apothecia were observed in June 2020 and only four plots were sampled in July. In 2021, the apothecia appeared mid-June and all the sites were sampled both in June and July. In 2012, 2016, 2017 and 2020, the observed density of infected rachis was significantly higher than 2021 with a mean between 4.3 and 8.7 g.m^2^ in forest and between 2.7 and 16.0 in hedge and small wood (some values over 20 g.m^−2^ in 2012), whereas the infected rachis density in 2021 was less than 5 g.m^2^ (1.3 ± 0.4 g.m^2^ in forest and 0.8 ± 0.2 g.m^2^ in hedges/small woods, p<0.001). The gradient of infected rachis obtained allowed us to highlight a dependence of the fruiting rate of *H. fraxineus* with the density of infected rachis on the litter (Fig. 6a). Indeed, the estimated Allee threshold was significantly different from 0, with very similar values in forest (1.9 IC [1-2.9] g.m^−2^) and in hedge and small wood (1.5 IC [0.8-2.5] g.m^−2^). Below the threshold, the fruiting rate increased significantly with the density of infected rachis (μ =93.8 IC [59.5-153.3] in forest and 63 IC [39.3-93.5] in hedge and small wood). For density values above the Allee threshold, the fruiting rate did no depend on the density of infected rachis with a mean fruiting rate given by the parameter **A** of the Gompertz equation. The mean fruiting rate was different according the environment with a higher value in forest (88.6 IC [52.2-124.8] apothecia.g^−1^) than in hedge and small wood (48.2 IC [30.3-65.8] apothecia.g^−1^). The parameter **λ** was no significantly different to 0 (0.01 IC [0-0.037]) (Fig.6a). The rachis infection rate to reach the Allee threshold depended on the amount of rachises produced by the ashes present in the stand, so on ash basal area (Fig.6b). The Allee threshold would be reached at an infection rate inferior at 0.2 for an ash basal area higher than 20 m^2^.ha^−1^ and need an infection rate higher than 0.5 for low ash densities (< 5 m^2^.ha^−1^). In 2020, 72 % of studied ash stands had an infection rate sufficient for a *H. fraxineus* inoculum production not subjected to the Allee effect, whereas only 27% of the ash stands exceeded the Allee threshold in 2021 (Fig. 6b).

**Fig. 6.**
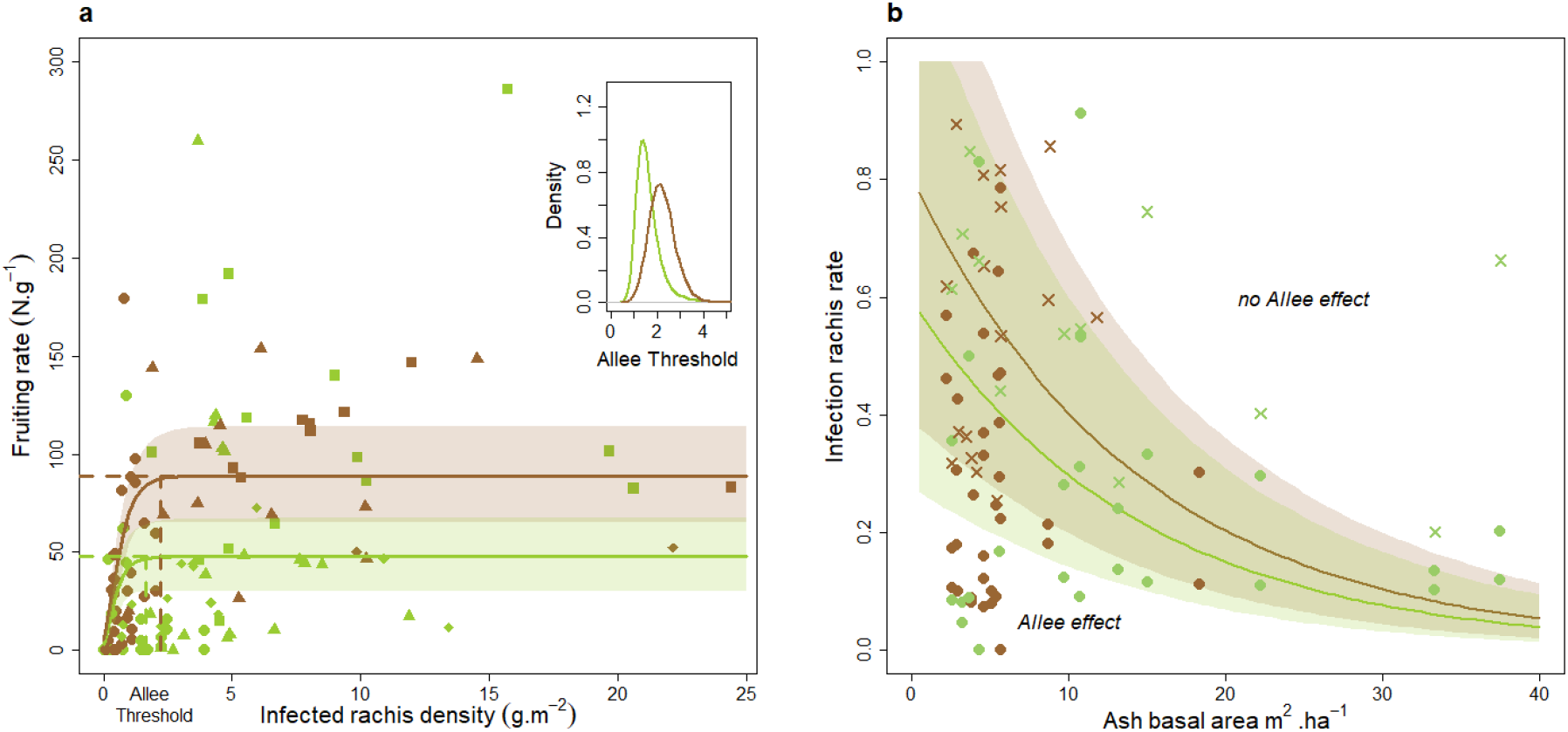
**a** Evolution of the fruiting rate as a function of the infected rachis density. In brown, measure from forest, in green, measure from hedge and small wood with their Gompertz function associated with year 2012 (diamonds), year 2016 (squares), year 2017 (triangles) and year 2021 (circles). **b** Estimated infection rate as a function of ash basal area to reach the Allee threshold in forest (brown curve) and in hedge and small wood (green curve), with measured infection rate in each situation in 2020 (cross) and 2021 (circle). Shaded areas correspond to 97.5% confidence intervals

The amount of ascospores detected in the spore traps in 2021 significantly increased with the density of apothecia observed in the plot litter in 2021 (0.01 p<0.05, Fig. 7a) and no differences were observed between forest and hedges/small woods (p>0.05). In addition, the quantity of ascospores detected in the spore traps in 2020 and 2021 was positively correlated with the density of infected rachis observed in the same year and also depended on plot environment and year (Fig. 7b). Indeed, the amount of trapped ascospores was higher in forests (0.26 in 2020 and 0.7 in 2021 p<0.01) than in hedge and small woods for similar infected rachis density (−0.03 in 2020 and 0.4 in 2021 p<0.01). Furthermore, the amount of ascospores trapped was significantly higher in 2021, than 2020 in particular in hedge and small wood where the amount of ascospores trapped was very weak (Fig. 7b).

**Fig. 7.**
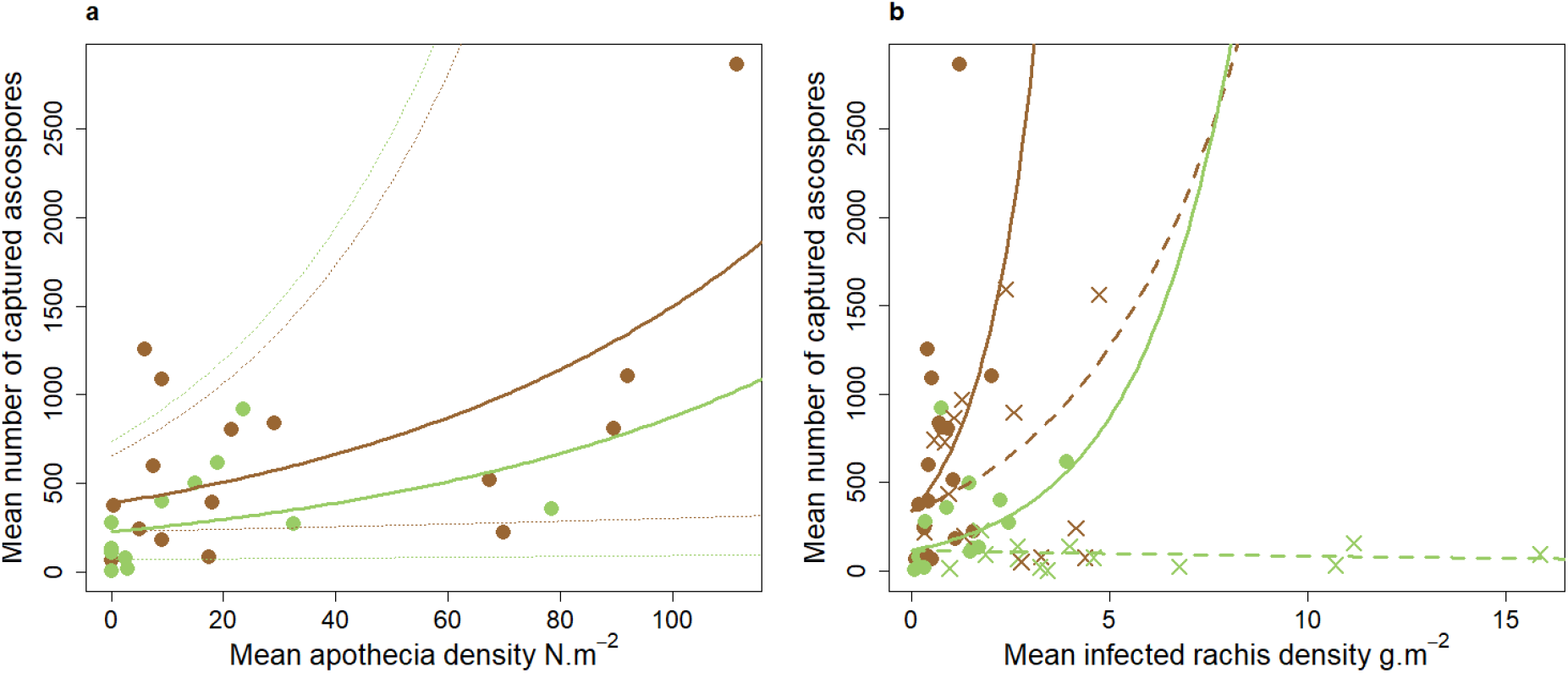
**a** Mean of ascospores captured in the ash stand according to the apothecia density for the year 2021. Brown points represent forest environment and green points represent the hedge and small wood environment. **b** Relation between mean of ascospores captured and infected rachis density in forest environment (brown) and hedge and small wood (green) during summer 2020 (cross points) and summer 2021 (circle points). Line represents the regression for 2021 and dashed line year 2020 (brown: forest, green: hedge and small wood)

## Discussion

This study analyses the relationship between ash, its health status and the population dynamics of *H. fraxineus* in relation to the stand environment. Our results revealed that host density was an important factor in the development of *H. fraxineus* populations, their inoculum production and subsequent health impacts on ash. We show that a component Allee effect on the fruiting rate exists for this fungal pathogen, confirming the hypothesis suggested by Hamelin et al. (2016), and that it limits inoculum production in the ash stands studied.

The larger negative impact of *H. fraxineus* on crown health at higher ash density that we observe is consistent with what has been reported in previous studies (Grosdidier et al. 2020; Havrdová et al. 2017), although this effect could not be observed early in the ash dieback epidemic (Bakys et al, 2013; Marçais et al, 2016). The average crown decline measured in the present study was in the same range as the one observed in 2016 and 2017 in the same area of Champenoux (Grosdidier et al. 2020): The value in forest was of 40% in 2016, 2017 and in 2020; and in none forest locations, slightly higher than 20% in the three study years of Grosdidier et al (2020), and 19% in 2020. This might suggest that ash health status stabilized within the past 5 years, so less than 10 years after ash dieback was first observed in Champenoux (2010, see Grosdidier et al, 2020). This seems surprising because mortality caused by ash dieback has been reported to develop late, approximately 10 years after the pathogen arrives in old trees (Marçais et al. 2017; Madsen et al. 2021) and it has been reported that ash mortality does not stabilize in the 15 first years of the epidemic (Coker et al. 2019).

Several features might explain that discrepancy. On the one hand, the ash density in the studied forest is low and decreased even more as the few pure ashes stand present were clear-cut for sanitary reasons between 2017 and 2020. Heavy decline and mortality occurred in the first decade of the epidemic and the logging of severely dieback trees probably removed the less tolerant ashes (Cleary et al. 2017; Børja et al. 2017; Skovsgaard et al. 2017). The remaining ashes might be more tolerant individuals which would limit dieback severity but not *H. fraxineus* presence. On the other hand, severe heat-waves and droughts periods occurred in the area in 2015, 2018, 2019 and 2020. The development of *H. fraxineus* is strongly limited by temperature above 35°C (Hauptman et al. 2013, Grosdidier et al. 2018). In addition, apothecia production and ascospore release are influenced by air and soil humidity (Dvorak et al. 2016; Gross et al. 2012; Hietala et al. 2013; Kirisits and Freinschlag 2012; Schumacher 2011). Therefore, infection rate of ash rachis in the litter and the crown decline may have been reduced by high temperatures in previous summers (Grosdidier et al. 2018). The drought observed in June and July 2020 may have prevented the apothecia production which would explain the low amount of ascospores that we observed in our spore traps in 2020 and the low proportion of rachis colonized by *H. fraxineus* in the litter 2021.

Grosdidier et al. (2020) reported that ash trees in hedges and small woods in the Champenoux area were healthier than those in forested areas and that the crown decline remained stable at about 20% from 2012 to 2018. Our data confirmed that the crown decline observed in 2020 and 2021 in hedge and small woods remained at about 20%. Furthermore, as noticed by Grosdidier et al (2020), we showed that, despite the greater decline of ash trees in forest conditions, the total amount and colonization rate by *H. fraxineus* of ash rachis in the litter were similar in hedges and small wood plots than in forest plots. Despite a similar infection rate of rachises, we observed lower apothecia production on them in hedges and small wood compared to forest locations in 2012, 2016, 2017 (Grosdidier et al, 2020) and 2021. This lower apothecia production in non-forested areas, certainly due to lower humidity level, may partly explain this impact difference of *H. fraxineus* on ash dieback in hedge and small wood although the infection rate of rachises does not differ between the two environments.

The demonstration that *H. fraxineus* is subjected to a component Allee effect linked to mating success offers a new insight into the invasion biology of the pathogen. Although the existence of this component Allee effect was suggested by (Hamelin et al. 2016), it has never been shown that it effectively limits inoculum production in ash stands. The Allee threshold was estimated to be of a similar magnitude in forests and in hedge and small woods (respectively, 1.9 IC [1-2.9] and 1.5 IC [0.8-2.5] g of infected rachis.m^−2^). The parameter **λ** was estimated to be near 0 which means that, even at very low rachis density in the litter, the mating success remains greater than zero. Thus, the observed effect can be considerate as a weak Allee effect. At low ash density, a strong infection rate is necessary to reach the Allee threshold. According our results, in an ash stand with a density lower than 5 m^2^.ha^−1^, the *H. fraxineus* population could be subjected to Allee effect below an infection rate of 50%. In years unfavourable to leaf infection, like 2020, such an infection rate was seldom reached.

Consequently, the development of ash dieback is highly dependent on ash density. At low ash density, significant apothecia and inoculum production occurs only when the infection rate of rachis is high. The component Allee effect might be expected to reduce inoculum production when infection is scattered in newly infected areas, and thus should reduce the overall dispersion rate (Lewis and Kareiva 1993; Taylor and Hastings 2005). This is consistent with the hypothesis raised by Hamelin et al (2016) and may explain why the dispersal speed observed in France has remained constant over time, *i*.*e*. around 60 km per year (Grosdidier 2017). This dispersal speed is very similar to what has been observed elsewhere in Europe (between 30 and 75 km per year Børja et al. 2017; Queloz et al. 2017; Ghelardini et al. 2017). This component Allee effect could also explain that the introduction of the pathogen through the planting of infected seedlings may result in foci that remain limited for an extended period of time. This was observed in central England, where dendrochronological analyses revealed the presence of *H. fraxineus* as early as 2005 in ash tree plantations remote from any other sources of inoculum, that is seven years before the pathogen was first reported in the country (Wylder et al. 2018).

In addition, the increase of the disease severity could lead to contain the pathogen population growth. Indeed, we observed, as expected, a positive relationship between total rachis density in the litter and ash basal area. However, the rachis production decreases with crown decline, the total rachis density of declining stands were strongly reduced compared to a healthier stand. As a consequence, the suitable reproduction substrate available for *H. fraxineus* becomes scarcer as the dieback progress. The density of infected rachis in the litter, which is a good measure of *H. fraxineus* population size, may also be reduced by the tree dieback. This could lead to infected rachis densities below the Allee threshold for mating success. A population density below the Allee threshold has a lower fruiting rate, the amount of ascospores release is weaker, the infection rate will decrease and this could lead local population extinction.

The last point concerns the potential impact of an Allee effect on the pathogen genetic diversity during the invasion process. Indeed, it was shown through modelling by Roques et al. (2012) that a population subjected to an Allee effect spreads as a pushed wave which results in a genetic diversity that remains stable throughout the colonization process. In this case, the bottleneck-induced loss of genetic diversity for a long-dispersal founder event is rare. Such founder events are usually characterized by low population density and strongly limit the growth of populations subject to the Allee effect (Roques et al. 2012). Noteworthy, Burokiene et al (2015) showed that *H. fraxineus* populations present in eastern Europe, in anciently colonized areas present similar genetic diversity compared to populations present on the expansion front. This is in line with the hypothesis of the invasion of Europe by pushed waves mechanism.

The management suggested to date to control ash dieback disease in affected stands is to thin stands to reduce host density (Skovsgaard et al. 2017; Enderle et al. 2019; Short and Hawe 2019). The known consequences were a drier and warmer microclimate due to reduced tree cover and openness to light. But, as we have shown, lower ash density also results in lower total rachis density in the litter and, as a consequence, in lower infected rachis density. However, inoculum production depends on this density of infected rachis, especially when it is inferior to the Allee threshold. Therefore, a reduction in rachis density by thinning ash trees will decrease inoculum production and the Allee effect component may exacerbate this mechanism. This kind of management of ash stands to reduce the production of *H. fraxineus* inoculum could be beneficial to the health of the ash trees. Indeed, the severity of damage is directly related to the level of inoculum present in the area. In case of high inoculum presence, the pathogen induces not only crown dieback but also collar necrosis, pathway to the establishment of other aggressors such as *Armillaria spp* (Husson et al. 2012; Madsen et al. 2021; Marçais et al. 2016).

Regarding the future of ash in Europe, it appears that after 10 years of the epidemic, some ash trees remain relatively healthy. As tolerant ash trees have been shown to produce more seeds than declining individuals (Semizer-Cuming et al. 2019), their offspring may be increasingly adapted to the disease, which should be good news for future stands in the region. Moreover, non-forest environment seems have some conditions unfavourable to the disease development, despite an infection rate similar to forest environment, which allows to ash to stay a structuring species of landscape within hedge and small wood.

## Acknowledgements

We wish to thank Olivier Caël and Arielle Beltran for his large involvement in data collection and Mireia Gomez-Gallego for here useful comments on the manuscript. The work was funded by the Homed H2020 project (grant no. 771271). The UMR1136 research unit is supported by a grant managed by the French National Research Agency (ANR) as part of the “*Investissements d’Avenir*” program (ANR-11-LABX-0002-01, Laboratory of Excellence ARBRE).

## Statements & Declarations

### Funding

The work was funded by the Homed H2020 project (grant no. 771271). The UMR1136 research unit is supported by a grant managed by the French National Research Agency (ANR) as part of the “Investissements d’Avenir” program (ANR-11-LABX-0002-01, Laboratory of Excellence ARBRE).

### Conflict of interest

The authors declare to have not conflict of interests.

### Author Contributions

The design of this study was developped by Benoit Marçais and Simon Laubray. The data were collected by Simon Laubray. The statistical analyses were performed by Benoît Marçais and Simon Laubray. The first draft of the manuscript was written by Simon Laubray and all author commented and corrected it. The final version was revised and agreed by all author.

## Notes

### Competing Interest Statement

The authors have declared no competing interest.

